# Genetic Association of *PLAG1* Gene Variant 14:25015640G>T with Wither-Height in Pakistani Cattle

**DOI:** 10.1101/2023.06.18.545456

**Authors:** Muneeba Naeem, Sarosh Zahid, Seemab Fatima, Muhammad Hassan Raza, Muhammad Osama Zafar, Rashid Saif

**Affiliations:** Department of Biotechnology, Qarshi University, Lahore, Pakistan; Decode Genomics, Punjab University Employees Housing Scheme, Lahore, Pakistan

**Keywords:** Cattle, body-size, *PLAG1*, Cattle stature, skeletal-frame, Mutation

## Abstract

The cow is one of the most valuable domesticated animals, producing milk, meat, fiber, hide and manure for agricultural purposes to serve humanity. Particularly, first two production traits are significantly linked with the physical characteristics of the animal e.g., wither-height, large body-size and skeletal frame are positively correlated with these production traits. The *PLAG1* is one of the many genes that has been associated with the aforementioned trait in many livestock and human species. So, the current study was conducted on a total of 50 cattles using ARMS-PCR genotyping technique followed by genetic association testing of the 14:25015640G>T variant using PLINK data analysis toolset. Our findings showed that 24% of the sampled Pakistani cow population are homozygous wild-type (GG), 12% homozygous mutant (TT), while 64% found heterozygous (GT). Sampled cows were obeying Hardy Weinberg Equilibrium (HWE) with *χ*2 (2, N = 50) = 10.39, *p* = 0.049. Similarly, Chi-square variant association was also observed significant with *p-*value of 1.267×10^−3^ having minor-allele frequency of 0.60 and 0.28 in heighted (cases) and control cohorts respectively. Additionally, a positive odds-ratio of 3.85 is also evident that the subject variant is under-selection and showing the tendency of the mutant allele almost 4-times higher in cases vs control groups. This pilot scale study would be helpful to gain insight of variability of the subject SNP in the sampled cow population but further functional studies on larger sample size may be conducted for validation and subsequent results can be disseminated to improve this valued trait of the indigenous cows for gaining maximum milk and meat production from this esteemed species.

## Introduction

Cattle have been an integral part of human history. Initially, our ancestors sought this animal for sustenance and many other purposes including leather provision and draught purposes. Over the ∼10,000 years, farmers have selectively bred and raised this animal particularly for enhanced meat and milk production [1]. As animal husbandry is one of the strong economic indicators in agriculture-based countries, particularly beef fattening, cattle is the most potential livestock species on a global scale. During the newnate calf judging process, the initial assessment parameter are birth weight (BW) [2] and calving ease which are the direct indicators of animal’s growth and its carcass potential [3]. Cattle meat is also important for our health due to high content of iron, mineral and essential vitamins e.g., niacin, riboflavin and vitamin K, B6/B12 which reduces tiredness, fatigue, provide proteins and enhance the muscles strength [4].

The interaction of geographical regions, animal species and specific breeds along with different rearing systems give rise to distinct challenges in ensuring animal security. These challenges can have significant implications for both animal and human public health [5]. Livestock plays a crucial role in ensuring food and nutrition security due to several factors, including the high nutritional value of animal-derived food products. The recent evolution of cattle is marked by fluctuation in its body-size. Height in the *Bos taurus* lineage was reduced by a factor of ∼1.5 from the Neolithic to the middle ages and expanded again only during the early modern ages [6]. Using heliotype analysis, it is evident that the bovine *PLAG1* mutation have major effects on body-size, weight and reproduction ∼1000 years old derive allele that increased rapidly in its frequency in northwestern European *B. taurus* between the 16-18^th^ centuries [7].

In livestock, cattle stature is highly influenced by the Pleomorphic Adenoma Gene 1 (*PLAG1*). The researcher first discovered this gene when they studied pleomorphic adenoma (PA) in human salivary glands that encodes zinc protein family, find in growth-related QNTs [8]. The PLAG1 gene that regulate various cellular processes. Hence, it has been identified as one of the players in controlling body stature such as the development of skeletal-frame and overall body-size in cattle and other species [9, 10]. According to this, *PLAG1* gene-specific variants can affect the activity or expression influencing skeletal muscle growth by regulating the production of growth factors and hormones that are involved in bone formation and development of height in cattle [6].

In the current study, the *PLAG1* gene variant 14:25015640G>T has been selected to genotype and to check its association with wither-height in Pakistani cattle population. Aformentioned *PLAG1* gene variant is located on Chr.14 ID AC_000171.1, transcript ID XM_005215432.2 (r.4098) and protein ID XP_005215489.1 in Bos_taurus_UMD_3.1.1 (GCF_000003055.6) assembly with rs109815800. Moreover, the stature of cattle contributes valuable insights for the advancement of more accurate genomic prediction models and to desiminate this variant in other cattle breeds and other livestock species as well upon the further functional validation studies.

## Materials and Methods

### Sample collection and DNA extraction

Blood samples of 50 cattle were collected to explore the association of *PLAG1* gene with height and skeletal stature of Pakistani cattle. For sampling, two groups were made, one is categorized as a heighted-cohort (n=25, wither-height >130cm at first parity) and the other one as a control-cohort (n=25, wither-height<130cm at first parity) (Figure 1). Blood samples were stored in K3-EDTA vacutainers and kept at 4°C for further usage. DNA was extracted from the sample using inorganic method as per standard protocol.

**Figure 1:**
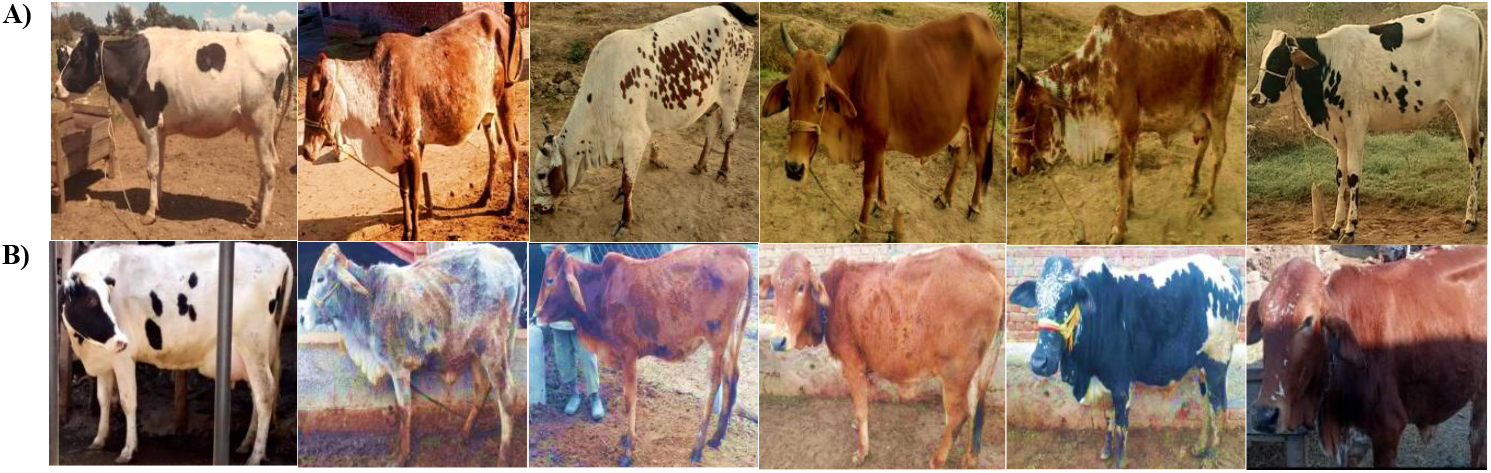
Few of the sampled cows, upper row animals are of whither-height >130cm at first parity **A)**, lower-row animals are below this threshold **B)**

### Primer designing for ARMS-PCR

The OligoCalc and NetPrimer softwares were used to design ARMS primers from the transcript ID XM_005215432.2. Five primers were designed, three of which are reverse normal (N), reverse mutant (M) and forward common. These primers were designed to amplify both wild and mutant-type alleles. The reverse wild/mutant ARMS primers also have a secondary mismatch at 3rd nucleotide position from 3’ end to increase the allele-specificity. As an internal control (IC), two additional primers (forward and reverse) were also designed to check the PCR fidelity. The detail of all the primers is provided in (Table 1).

**Table 1:**
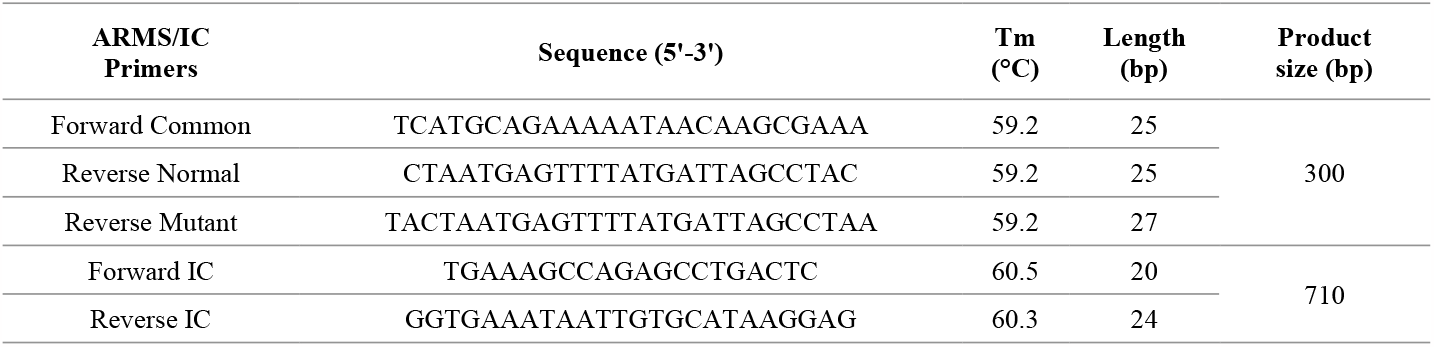
ARMS-PCR primers sequences details

### DNA amplification

An Applied Biosystems SimpliAmp thermal cycler was used to perform the ARMS-PCR reaction. The normal (N) and mutant (M) type ARMS allele-specific reverse primers and a common forward primer were used in two independent PCR reactions alongwith two regular primers to amplify the genomic region as an internal control. The reaction mixture of total 12μL volume contains 1μL of 50ng/μL genomic DNA, 10mM of each primer, 0.05IU of *Taq* polymerase, 2.5mM MgCl2, 2.5mM dNTPs, 1x buffer and molecular grade water. Five minute initial denaturation at 95°C was followed by 30 cycles of denaturation at 95°C for 45 sec, annealing at 60°C for 30 sec and extension at 72°C for 45 sec with the final extension at 72°C for 10 mins followed by storage of 4°C for infinity.

### Statistical analysis

Hardy Weinberg Equilibrium (HWE) equation *p*2 + 2*pq* + *q*2 = 1 was applied to check whether our sampled population obeying the above equation or not, followed by Chi-square analysis 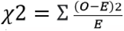 equation after determining the genotypic and allelic frequencies to evaluate the statistical association *p*-value. Moreover, the odd-ratio was also calculated using PLINK data analysis toolset.

## Results

In the current study, *PLAG1* variant 14:25015640G>T (rs109815800) was genotyped and found variable in Pakistani cattle population. A total of 50 samples were genotyped (n=25 with wither height ≥130cm, and n=25 with wither height <130cm) at first parity. After statistical analysis, we found 12 animals are homozygous-wild type (G/G), 6 homozygous-mutant (T/T) and 32 heterozygous (G/T) in the overall sampled population. While in control group 11 cattle were homozygous-wild, 14 were heterozygous while none of the animal was observed as homozygous mutant. Similarly in cases 01, 18 and 06 cattle are homozygous wild-type, heterozygous and homozygous-mutant respectively as shown in (Figure 2).

**Figure 2:**
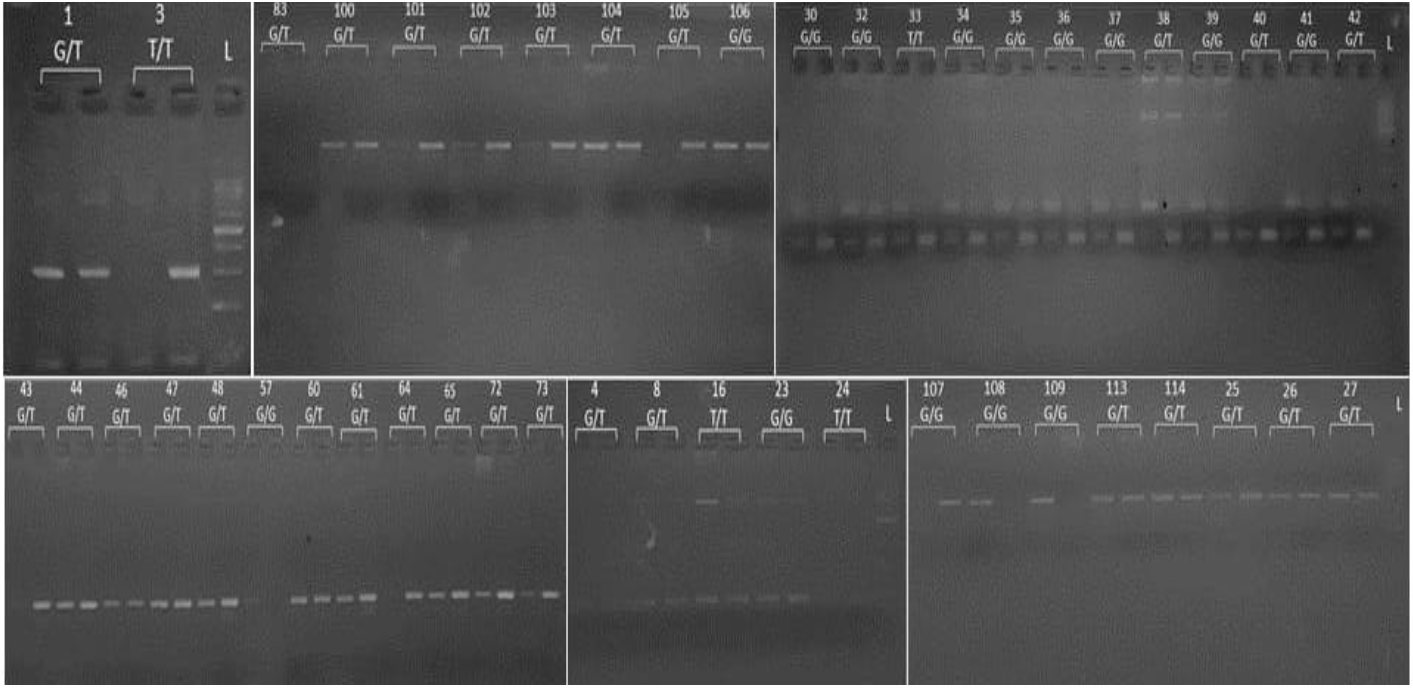
ARMS-PCR amplification of targeted variant within sampled cows, internal control and allele-specific product size of 700 & 300bp respectively.

Subsequently, HWE Chi-square analysis was applied to sampled population having *p-*value = 0.049 that is above the set threshold confidence interval which exhibit the obedience of HWE. Furthermore, genetic association was also infered which gave mutant allele frequencies of 0.60 in cases and 0.28 in controls having *p*-value 1.267×10^−3^ showing its significant association with body-height phenotype in Pakistani cattle. Similarly, odds-ratio (OR) of 3.85 was observed which expresses the prevalence of odds/mutants is almost 4-times higher in cases versus controls as described in (Table 2).

**Table 2:** Statistical association of subject variant with Pakistani cows wither-height

| No. of samples | Chr. Position | cDNA variant<br>XM_005215432.2 | Protein variant<br>XP_005215489.1 | Genotypic information | | | Alternative allele frequency | | $p$-value & (OR) |
| --- | --- | --- | --- | --- | --- | --- | --- | --- | --- |
|  |  |  |  | Wild type<br>GG/% | Hetero<br>GT/% | Mutant type<br>TT/% | Cases | Control |  |
| 50 | AC_000171.1<br>14:25015640 | rs109815800<br>(G>T) | p.= | 12/22 | 32/64 | 6/12 | 0.6 | 0.28 | $1.267 \times 10^{-3}$<br>(3.85) |

The multiple sequence alignment of *PLAG1* gene was also performed to find out the conservancy status of the genotyped variant in total 13 mammalian species of pig, cattle, deer, donkey, horse, monkey, human, chimpanzee, tiger, wolf, beer, dog and fox. The high level of conservancy of the subject variant was found in all species except the tiger (Figure 3).

**Figure 3:**
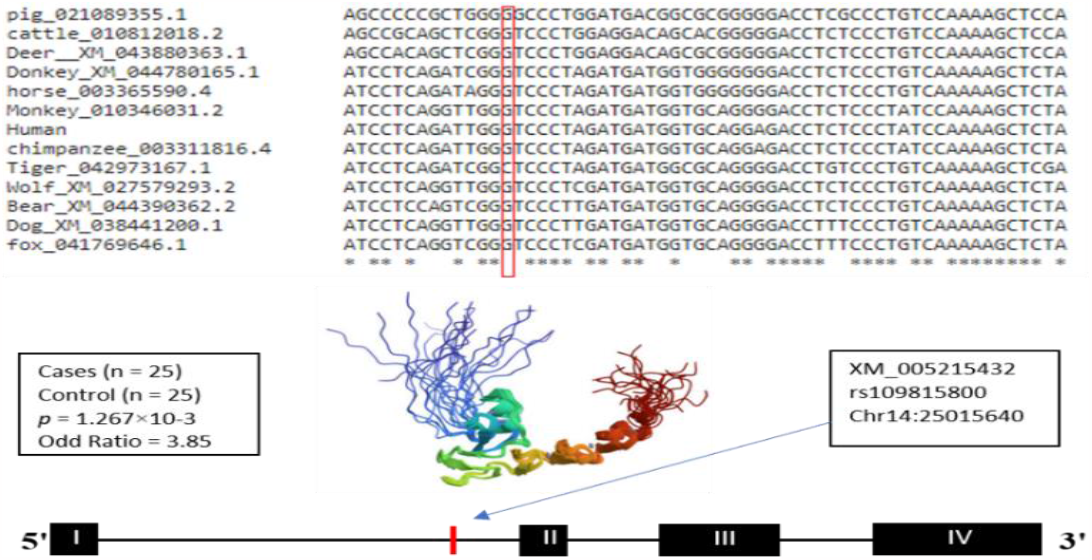
Multiple sequence alignment of genotyped variant of different mammalian species and genomic location.

## Discussion

In the current study cattle are classified into two groups based on their wither-height at first parity. The height of an animal at specific age is a advantageous indicator for determining its future growth and milk production potential due to more weight and blood circulation in the udder. Additionally, skeletal-frame can assist in predicting the fattening patterns of cattle breeding stock. However, the most accurate prediction of weight and performance in offspring resulting from breeding stock is obtained by estimated breeding values (EBVs) based on weight records [8]. Animal-height has also been found to be an informative indicator of performance in the feedlot, serving as the measure of maturity. Generally, larger animals tend to grow faster and accumulate less fat as compared to smaller ones. To discuss and evaluate height, it is advisable to use frame scores based on hip-height at specific age. Frame score is a generally accepted method for expressing the skeletal size of an animal. Most animals should maintain the same frame score throughout their lives, even as their actual height increases with age. Body type scores, ranging from 1 to 11 was developed at the University of Wisconsin-USA and applicable to almost all cattle breeds with little geographical variations [9].

The present study provides evidence suggesting that the selective pressure on the *PLAG1* gene may be traced back to recovery in cattle stature. Geographical distribution of a specific mutation exhibits an opposite pattern to that of stature, thus supporting its involvement in stature recovery [11]. Current study aim was to investigate the impact of genetic variation on the target attribute in the population of Pakistani cattle. The results revealed that there were 12 cows with the homozygous-wild type, 6 are homozygous mutant and 32 were heterozygous in both the cases and control groups, which support the aforementioned point of under-selection phenomenon. Additionally, multiple sequence alignment of the subject locus was conducted across 13 mammalian species to assess its conservancy status.

Previous researches has also been demonstrated that the *PLAG1* gene plays a key role not only in stature regulation in cattle but also in several other animal species as well [12]. Recent advancements in the field of stature genetics revealed that livestock possessed a relatively simple genetic architecture compared to humans. The *PLAG1* gene known for its involvement in adult human height has also been associated with selective sweeps in dogs and pigs as well as correlations between height and body weight in cattle and horses [13, 14]. These findings highlighted the significant impact of subjected genetic variant on mammalian growth and body size, underscoring the value of natural variants for selective breeding purposes. So, the current pilot scale study also provides another evidence that the same gene variant is significantly associated with the wither-height in Pakistani cattle too which is in conformity to the other livestock species of the world.

## Conclusion

It is being reported that *PLAG1* gene variant (rs109815800) is remarkably associated with the wither-height in Pakistani cattle population with *p* = 1.267×10^−3^. To increase the height and stature of the cattle herds for more meat and milk production, marker-assisted breeding policies may be employed by validating this variant after further functional studies.

## Ethical Statement

Cattle included in this study were privately owned, their blood samples were obtained from the farmers of personal acquintances upon their consent and collected under the supervision of veterinarians.

## Acknowledgements

Authors are obliged to the farmers for providing blood samples of their animals for this pilot scale study.

